# Expanding risk predictions of pesticide resistance evolution in arthropod pests with a proxy for selection pressure

**DOI:** 10.1101/2022.04.18.488707

**Authors:** Joshua A. Thia, James Maino, Alicia Kelly, Ary A. Hoffmann, Paul A. Umina

**Affiliations:** Bio21 Institute, School of BioSciences, University of Melbourne, Parkville, VIC, Australia; Cesar Australia, Brunswick, VIC, Australia

## Abstract

Chemical resistance in pest organisms threatens global food security and human health, yet resistance issues are mostly dealt with reactively. Predictive models of resistance risk provide an avenue for field practitioners to implement proactive pest management but require knowledge of factors that drive resistance evolution. Despite the importance of chemical selection pressure on resistance evolution, poor availability of chemical usage data has limited the use of a general multi-species measure of selection pressure in predictive models. We demonstrate the use of pesticide product registrations as a predictor of resistance status and potential proxy of chemical selection pressure. Pesticide product registrations were obtained for 427 USA and 209 Australian agricultural arthropod pests, for 42 and 39 chemical Mode of Action (MoA) groups, respectively. We constructed Bayesian logistic regressions of resistance status as a function of the number of pesticide product registrations and two ecological traits, phagy and voltinism. Our models were well-supported with demonstrated power to discriminate between resistant and susceptible observations in both USA and Australian species sets using cross-validation. Importantly, we observed strong support for a positive association between pesticide products and resistance status. Our work expands the horizon for proactive management by quantitatively linking a proxy for selection pressure on pest species to different chemical MoAs. This proxy for selection pressure can be combined with ecological information to predict the resistance risk in agricultural pests. Because pesticide product registrations can typically be derived from publicly available data, we believe there is broad applicability to other agricultural pests such as weeds and fungi, and to other geographical regions beyond the USA and Australia.

## BACKGROUND

Decades of research into the evolution of pesticide resistance and its molecular basis have contributed to a rich understanding of the dynamics of resistance evolution, which has benefited management strategies in agricultural and human health contexts (Crow 1957; Georghiou and Taylor 1986; McKenzie and Batterham 1994; Hawkins et al. 2019). But despite our understanding of pesticide resistance evolution, resistance remains a prevalent global issue for pest management and food security (Pimentel et al. 1992; Liu 2015; Sparks and Nauen 2015; Gould et al. 2018; Pu et al. 2020). Indeed, notwithstanding the best efforts of scientists, field practitioners and multi-national chemical companies, entire chemical Mode of Action (MoA) groups have become ineffective against certain pest groups due to widespread resistance (Nauen 2007; Bass et al. 2015; Horowitz et al. 2020).

A major challenge in managing pesticide resistance is the ongoing importance of chemical control, which places populations under considerable selective pressure to evolve resistance. Pest management strategies are often *reactive*, only being implemented once resistance has evolved and resistant genotypes have reached a detectable frequency in field populations (Tabashnik et al. 2014; Sparks and Nauen 2015; Umina et al. 2019). More *proactive* strategies would routinely assess resistance risks prior to control failures developing. Predictions about where and when a species might exhibit high risks of resistance evolution could be used to prioritize research investment and target in-field monitoring efforts. To achieve this, a comprehensive understanding of factors that increase the risk of resistance evolution is required.

There have been numerous efforts over many years to develop frameworks for inferring resistance risk as a combination of eco-evolutionary characteristics of pests and operational factors in agriculture. These include conceptual models (Georghiou and Taylor 1977, 1986; Audsley and Down 2015), decision trees based on qualitative factors (Rotteveel et al. 1997), and quantitative summarization of global resistance patterns in different species and chemical MoAs (Moss et al. 2019). More mechanistic insights have come from attempts to model covariates of resistance that incorporate ecological traits, structural characteristics of pesticides, and phylogenetic constraints in different pest groups (Rosenheim and Tabashnik 1991; Grimmer et al. 2015; Brevik et al. 2018; Hardy et al. 2018; Rane et al. 2019; Crossley et al. 2020). However, one of the most important risk factors of pesticide resistance evolution is selection pressure (Crow 1957; Georghiou and Taylor 1977; McKenzie and Batterham 1994). Yet quantifying the magnitude of selection pressure experienced by pest populations is challenging in agro-ecosystems because many countries lack policies for reporting pesticide usage, making regulation and risk assessment difficult (Handford et al. 2015; Sharma et al. 2019).

Studies investing in the co-collection of pesticide usage and phenotypic resistance data have provided valuable demonstrations of the link between on-farm practice and the evolution of pesticide resistance. For example, studies of weeds from the United Kingdom show that the spatial distribution of herbicide resistance is linked to historical application rates and frequency of herbicide applications (Evans et al. 2016; Hicks et al. 2018). Yet collecting such data requires considerable investment, especially across large spatial scales, and is limited in the scope of species and pesticides that can be assessed simultaneously. Despite limited availability of accessible databases that directly record pesticide usage, other data sources may provide information that can be used to construct covariates of pesticide usage. In prior work, we have demonstrated that registrations of pesticide products are correlated with pesticide usage in agricultural weed, fungal and arthropod pests from the USA and Australia (Maino et al. 2023). These findings suggest pesticide product registrations may be a general proxy for inferring selection pressure and the risk of pesticide resistance evolution across different pest groups.

In this present work, we directly test the ability for pesticide product registrations to act as a predictor of pesticide resistance status. We focus specifically on arthropod pests (insects and mites) in agro-ecosystems from the USA and Australia as a proof of concept. In line with our expectations, we demonstrate a positive association between pesticide product registrations and pesticide resistance status in agricultural arthropod pests from both countries. Our results suggest that pesticide product registrations can serve as a generalizable proxy for selection pressure in agricultural systems. Pesticide product registrations may therefore provide a useful quantitative measure for assessing resistance risk in different geographic regions for specific pest species and chemical MoAs (particularly when combined with ecological information), helping guide proactive resistance management efforts.

## METHODS

### Dataset compilation

Data on pesticide products, phagy and voltinism were collected for agricultural arthropod pests from the USA (*n* = 427 species) and Australia (*n* = 209 species). Our working dataset was compiled from publicly available databases and published studies. All data filtering was performed in R v4.1.3 (R Core Team 2022).

### Pesticide products

In the absence of detailed information on chemical usage patterns against agricultural pests in most parts of the world, our study takes the approach of using pesticide product registration data as a potential proxy for chemical usage and selection pressure. Our rationale is that economically important arthropod pests will have many pesticide products registered against them and will be exposed to greater pesticide usage in the field. Here, we strictly refer to “pesticide product registrations” as all unique product registrations associated with a specific chemical MoA group.

For USA pesticide products, we downloaded pesticide product registration data from the Environmental Protection Agency (EPA; https://www.epa.gov/; accessed February 11^th^, 2021). Likewise, registrations of Australian pesticide products were obtained from the Australian Pesticides and Veterinary Medicines Authority (APVMA; https://apvma.gov.au/; accessed February 9^th^, 2021). Each of these databases contain tens of thousands of product registrations across time that include agricultural and non-agricultural uses (EPA, *n* = 165,661; APVMA, *n* = 22,090). Keyword searches were used to isolate those registrations related to pesticide products for agricultural pests. For the EPA database, we filtered the “Site” variable for keywords such as “FOLIAR TREATMENT” and “SEEDLING” (See List S1 for full details). For the APVMA database, we filtered the “Category” variable for “AGRICULTURAL” products and then filtered the “Product_type” variable for the keywords: “INSECTICIDE”, “MITICIDE” and “MOLLUSCICIDE”.

Pest common names were assigned to species scientific names, and multiple common names per species were merged. For logistical reasons, we only considered the top 50% most registered species. First, the number of pesticide product registrations were pooled across MoAs within a pest common name to get the cumulative number of pesticide product registrations. Second, pest common names were ranked from most to least cumulative pesticide product registrations. Third, only those pest common names above the 50^th^ percentile were selected for downstream analyses. This procedure removed unique pest common names with very few pesticide product registrations and eased manual curation of our dataset. Taxonomic assignment of pest common names was initially automated with the R package *taxize* (Chamberlain and Szöcs 2013;

Chamberlain et al. 2020) and were manually assigned otherwise. Species lists were then compared against reference databases of species occurrence to remove any putatively incorrect species assignments. USA species were compared against a published inventory of USA pests (Hardy et al. 2018) and Australian species were compared against occurrence records in the Atlas of Living Australia (https://www.ala.org.au/; accessed July 7^th^, 2021). We also removed pests like cockroaches, invasive ants, or pests of ornamentals, conifers, and turf. We were left with 427 USA species (750 common names) and 209 Australian species (247 common names) that were agricultural pests.

Chemical MoA groups were assigned to each product registration based on their listed active ingredients from the Insecticide Resistance Action Committee (IRAC) classifications (Table S1). Next, we summed the number of unique chemical registrations per species for each chemical MoA across relevant pest common names. In both chemical databases, some chemical registrations are listed at higher order taxonomic classifications (*i.e*., genus or higher), for example: “aphids”, “thrips”, or “moths”. We considered two alternate ways to deal with these ambiguous assignments. First, all entries that cannot be assigned to species level could be removed. However, this would substantially decrease the number of entries in certain pests where many registrations are not at the species level (e.g., cereal aphids). Second, entries could be assigned to all species within that taxonomic rank. However, this would inflate the total number of entries for pests that are not targeted by a pesticide product but occur in the same taxonomic rank as a targeted species. This is expected to increase the similarity between targeted and non-targeted species with respect to the number of products registered against them, and hence reduce signals of selection pressure. To retain as much data as possible, we opted for the second approach, allocating pesticide products up to the family level, ignoring registrations at the order level or higher. Additionally, chemical mixtures provided a unique count for each chemical MoA present in the product. For example, a product containing both a pyrethroid (group 3A) and a neonicotinoid (group 4A) would comprise a pesticide product for each of these chemical MoAs separately.

### Phagy and voltinism

We collected data on two ecological traits previously associated with increased rates of pesticide resistance evolution: phagy (the number of host plant families) and voltinism (the number of generations per year). We leveraged the data published in Hardy et al. (2018), which expanded work presented in Rosenheim et al. (1996). For each candidate species, we extracted available values of phagy and voltinism. For those species not included in Hardy et al. (2018), we performed a literature search to acquire this information using Google Scholar (see Lists S2 & S3 for full details).

We were unable to collect trait data for every species. Priority was given to trait values of species in the Australian dataset, which were used for corresponding species in the USA dataset. We then used taxonomically focused imputation to obtain values for those remaining species missing trait data. For a given species with a missing phagy or voltinism value, an imputed value was derived from the mean of other congeneric species. If congeneric species were not present, or lacked data, the mean of confamilial species was used. Where it was not possible to obtain a mean from confamilial species, the mean of species within the same order was taken. This approach is conservative as it reduces the variance among species with shared taxonomy, although we demonstrate it has little effect on our general conclusions (see Results).

### Resistance status

For each species and chemical MoA group combination, we assigned resistance status (present or absent), as reported in the Arthropod Pesticide Resistance Database (APRD: Mota-Sanchez and Wise 2021) and from the literature. For Australian species, we combined APRD records with a literature search. For USA species, APRD records alone were used to assign resistance status. APRD records for both countries were filtered to only include those marked as “Field Evolved Resistances” to ensure we did not include any records of experimentally evolved resistance.

### Statistical analyses

#### Predictive models of resistance

We used Bayesian logistic regression to model resistance status in each species against different chemical MoA groups. Our ‘full’ model was constructed as follows:

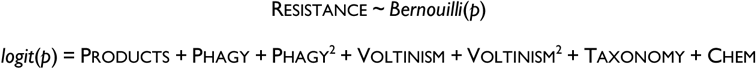

Resistance was a Bernoulli response for resistance status, 0 for susceptible and 1 for resistant, for each species to a given chemical MoA. Products (number of product registrations), Phagy (number of host plant families), and Voltinism (number of generations per year) were continuous fixed effects. Each of these fixed effect predictors were log-scaled: where 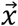 was the vector of predictor values, we applied the transformation log_10_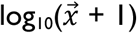 and then scaled to a mean of 0 and variance of 1. Because prior studies have indicated curvilinear relationships between resistance evolution and these two ecological traits (Rosenheim et al. 1996; Hardy et al. 2018; Crossley et al. 2020), we fitted first and second order orthogonal polynomials for Phagy and Voltinism using R’s *poly* package. Taxonomy was a random effect accounting for taxonomic structure, fitting a random intercept for each genus, nested in family, nested in order, as used elsewhere (Hardy et al. 2018; Crossley et al. 2020). CHEM was a random effect, fitting a random intercept for each chemical MoA group.

We fitted our ‘full’ model to pest datasets with imputed and non-imputed ecological traits to evaluate the effect of trait imputation on predictions. To evaluate the performance of our resistance predictors, we constructed a ‘null’ model that fitted the global mean and the random effects of taxonomy and chemical MoA group. A ‘products’ model was also fitted to test the predictive power of pesticide product registrations in isolation. All three models were fitted separately on the USA and Australian pest species sets in a Bayesian framework with R’s *brms* package (Bürkner 2017). All models were fitted using 10 chains, each with 8,000 iterations, for a total of 80,000 iterations. The first 50% of iterations were used as burn-in, leaving 40,000 iterations for estimating model coefficients. An adaptive delta value of 0.999 was used to set a high average target acceptance rate and prevent divergent transitions after the burn-in phase. The ‘full’ and ‘products’ models were compared against the ‘null’ model using the *bayestestR* R package (Makowski et al. 2019) to obtain Bayes factors. A Bayes factor >3, or a log-Bayes factor >1.1, indicated substantial support for a focal model relative to the null model (Wetzels et al. 2011).

#### Model generality

We used *K*-fold cross-validation to evaluate the generality and predictive power of our ‘full’ model and the ‘products’ model. We analyzed 100 training–testing partitions of both models used imputed datasets. For each partition, 85% of the resistant and susceptible species were randomly selected for the training subset, and the remaining 15% of resistant and susceptible species were allocated to the testing subset. Thus, the proportion of resistant to susceptible species was retained in both the training and testing subsets. The ‘full’ and ‘products’ models were fitted to each training subset using the *brms* function with 10 chains, 1,000 iterations per chain, and a burn-in of 5,000 chains, with 5,000 iterations for coefficient estimation, and an adaptive delta of 0.999. Predictions on the testing subsets were made using the *posterior_predict* function from the *brms* package without inclusion of random effects. Random effects were excluded because random subsampling of species led to non-overlapping combinations of order, family, and genus between the training and testing subsets. Because predictions were made without consideration of random intercepts for taxonomy and chemical MoA group, our cross-validation analyses provided a more conservative assessment of the generality of our predictive models. For direct comparison, ‘full’ and ‘products’ models were fitted to the same training–testing species subsets.

To quantify predictive power, we calculated AUC (area under the receiver–operator characteristic curve) scores for each testing subset. AUC scores describe the relative rate of true to false positives for increasing predicted risk scores. AUC scores range from 0–1; as values approach 1, there is increasing likelihood that resistant species will be assigned higher predicted risk scores than susceptible species. For each testing subset, observed (0 or 1) and predicted (probability from 0 to 1) values were passed to the *auc* function from R’s *Metrics* package to obtain an AUC score.

## RESULTS

### Dataset compilation

We compiled two datasets comprising agricultural arthropod pests from the USA and Australia, their ecological traits (phagy and voltinism), the number of pesticide products registered against them for different chemical MoAs, and their resistance statuses to those chemical MoAs. Our USA species set included 427 species, 81 families, and 10 orders, and for the Australian species set, there were 209 species, 65 families, and 12 orders. The taxonomic composition of total species was analogous between countries (Fig. 1). In the USA, Lepidoptera (32%), Hemiptera (29%), Coleoptera (19%), and mites from the order Trombidiformes (9%) made up the bulk of the total pest species, which was very similar to the Australian species set (30%, 29%, 18% and 9%, respectively). The taxonomic composition of resistant species (those with resistance to at least one chemical MoA) largely paralleled the total species set (Fig 1). In the USA, Lepidoptera (32%), Coleoptera (26%), Hemiptera (25%) and Trombidiformes (9%) made up the bulk of resistant pest species (*n* = 76). In Australia, resistant pests (*n* = 24) included species from Lepidoptera (46%), Hemiptera (21%), Trombidiformes (17%), Thysanoptera (8%) and Coleoptera (8%).

**Fig. 1.**
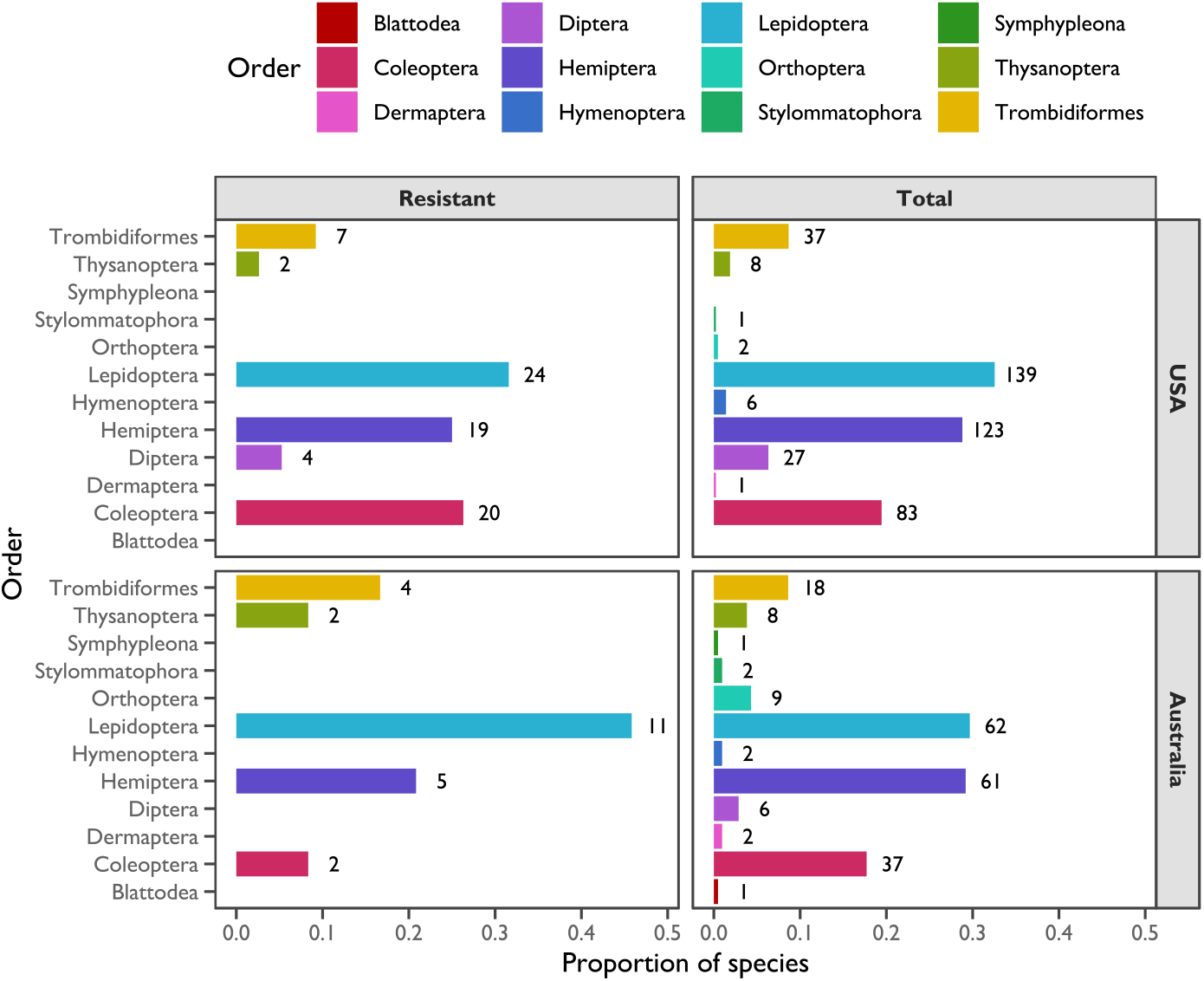
Taxonomic distribution of analysed agricultural arthropod pests. Panels illustrate the proportion of species from different taxonomic groups with resistance (left column) and the total species set (right column) in the USA (top panels) and Australia (bottom panels). Taxonomic orders are on the *y*-axis, and the proportion of species is on the *x*-axis. The length of bars indicates the proportion of species, coloured by order (see legend), with numbers to the right of bars noting the number of species.

Values of phagy and voltinism were sourced from previous studies (Rosenheim et al. 1996; Hardy et al. 2018; Crossley et al. 2020) or were sourced from a literature search. However, many species required imputation. Of the USA species, 30.9% (*n* = 132) had at least one ecological trait imputed from a literature search, whereas for Australian species, 56.0% (*n* = 117) of species had at least one ecological trait imputed. The USA species set contained 42 chemical MoAs and the Australian species set contained 39 chemical MoAs. Our working datasets were structured with respect to species–MoA observations, where the number of pesticide product registrations for each unique chemical MoA was assigned to each pest species. There were *n* = 17,934 species–chemical MoA observations in the USA species set and *n* = 8,151 in the Australian species set.

### Predictive models of resistance

Our ‘full’ model of resistance fitted resistance status of species to different chemical MoAs (a binary response) as a function of the fixed effects of pesticide product registrations (*β*_Products_), the first and second polynomials of phagy (*β*_Phagy^1_, *β*_Phagy^2_) and voltinism (*β*_Voltinism^1_, *β*_Voltinism^2_), and the random effects of taxonomy and chemical MoA. We also fitted a ‘products’ model that only included the fixed effect of products and the random effects of taxonomy and chemical MoA, and a ‘null’ model that only fitted the random effects of taxonomy and chemical MoA. Our models were fitted using a Bayesian framework and the likelihood of the ‘full’ and ‘products’ models were tested by comparison to the ‘null’ model.

We tested whether ecological trait imputation affected the predictions of the ‘full’ model by comparing models using imputed values of phagy and voltinism (all species) with models using non-imputed values (excluding species with missing data). Because predicted risk scores using the ‘full’ model using imputed and non-imputed ecological traits exhibited very high correlations (*r* > 0.98; Fig. S1), all our interpretations consider those models fitted to datasets with imputed ecological traits. Distributions for fixed effects are illustrated in Fig. 2. Fitted terms for the ‘full’ models are tabulated in Table 1 and illustrated in Fig. 2.

**Fig. 2.**
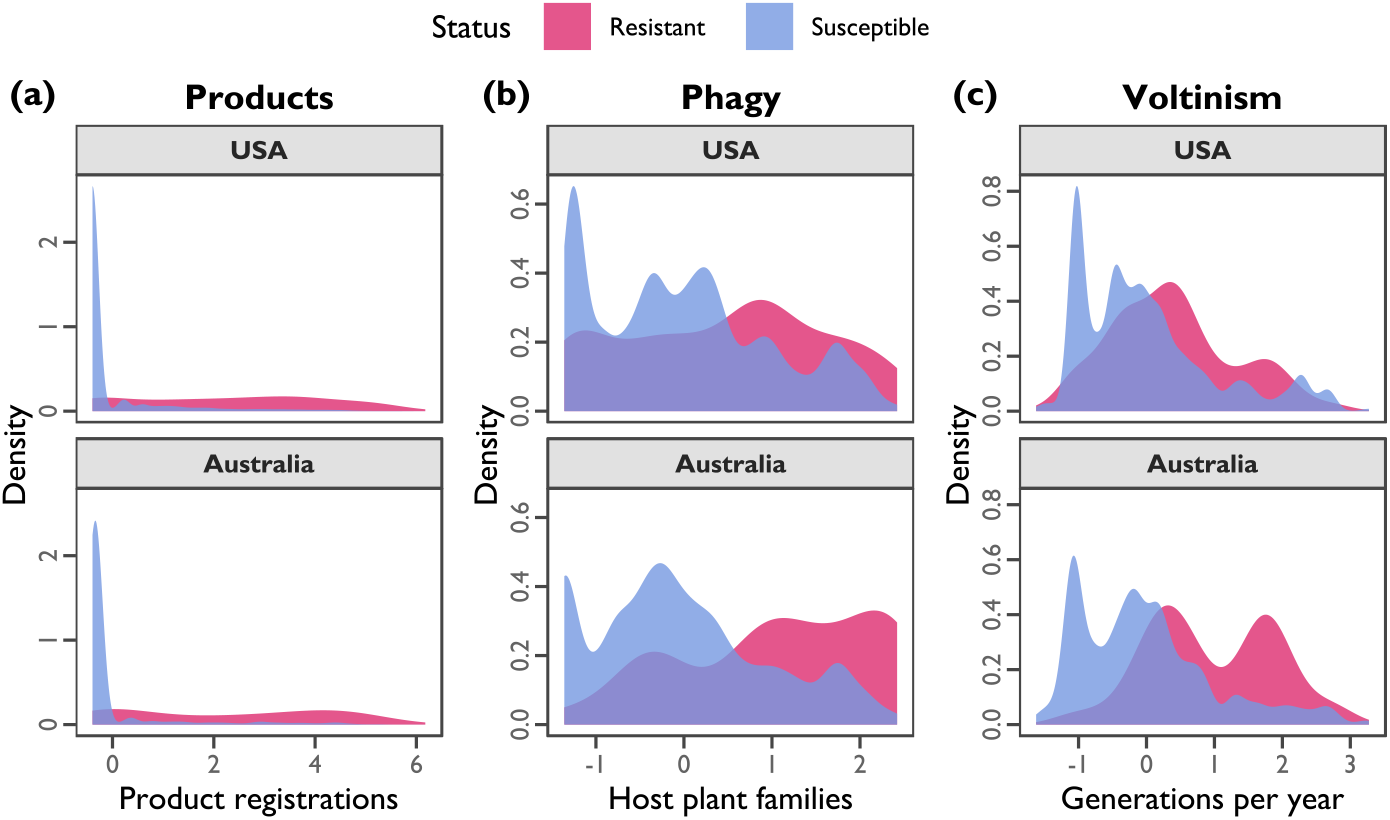
Resistance predictor distributions. Curves depict the kernel density estimate, with area under the curve (AUC) equal to 1.00. Resistant species–chemical MoA observations are depicted by pink curves and blue curves depict susceptible species–chemical MoA observations. Panels contain observations from the USA and the Australian species sets separately. (a) Products, the log-scaled number of product registrations. (b) Phagy, the log-scaled number of host plant families. (c) Voltinism, the log-scaled number of generations per year.

**Table 1.**
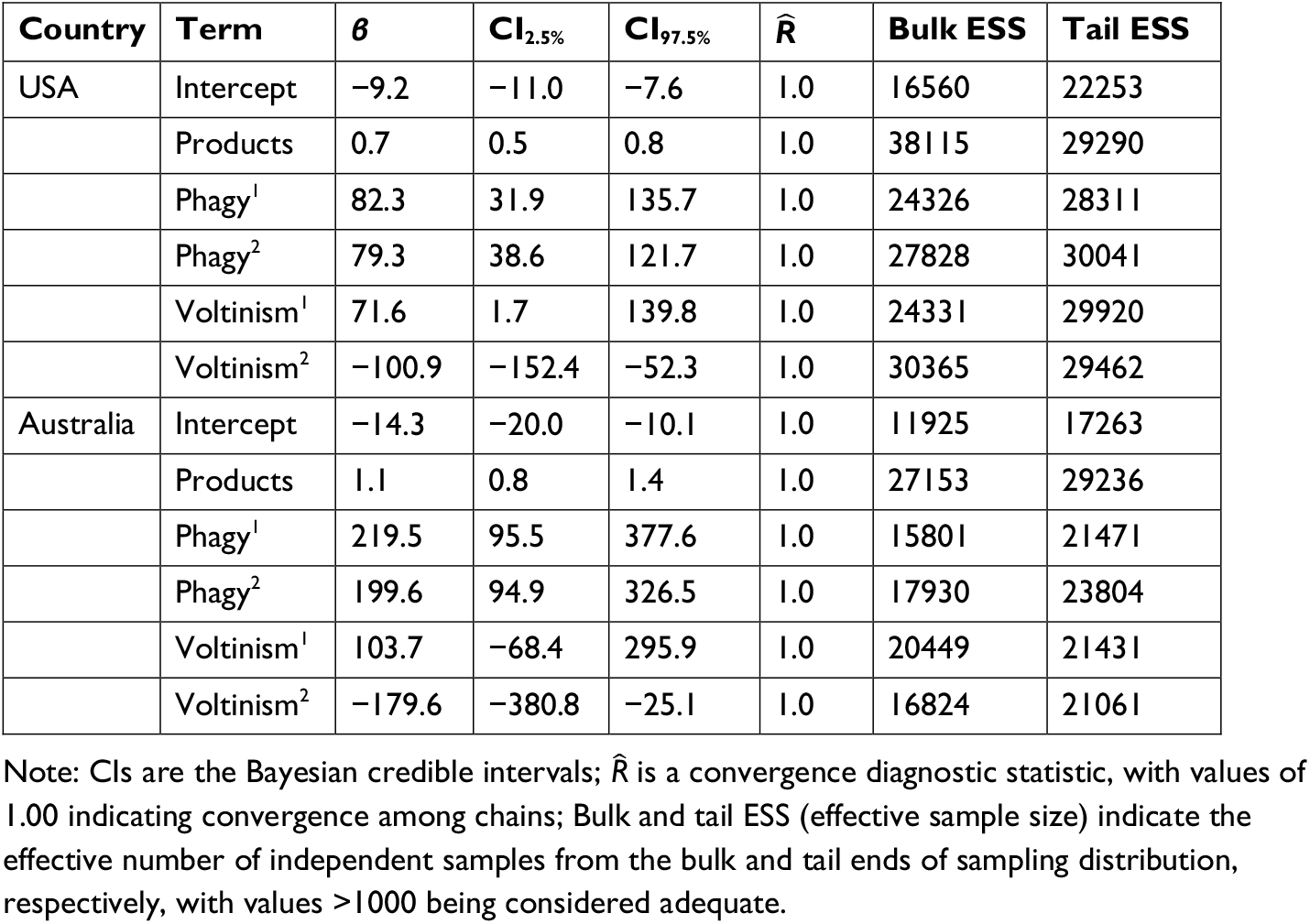
Posterior estimates of model terms and Bayesian diagnostic statistics for ‘full’ models of resistance

We found a positive association between pesticide product registrations and resistance status (Figs. 2 & 3). For a given chemical MoA, species with a larger number of pesticide product registrations were more likely to be resistant to that chemical MoA than those with fewer pesticide product registrations. This effect was consistent between species sets from the USA, *β*_Products_ = 0.68, and Australia, *β*_Products_ = 1.09, with overlapping Bayesian 95% credible intervals (Table 1; Fig 3). Consistent with other studies, we found a positive second order polynomial for phagy and a negative second order polynomial for voltinism, suggesting that resistance evolution is minimal at intermediate levels of phagy and maximised at intermediate levels of voltinism (see also Hardy et al. 2018; Crossley et al. 2020). We observed consistent estimates in the second order polynomials for both countries, with overlapping Bayesian 95% credible intervals: for USA species, *β*_Phagy^2_ = 79.30 and *β*_Voltinism^2_ = −100.90; and for Australian species, *β*_Phagy^2_ = 199.58 and *β*_Voltinism^2_ = −179.64 (Table 1; Fig. 3). Diagnostic statistics indicated that we reliably sampled the posterior distributions for these fixed effect coefficients (Table 1); the 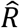 statistics were equal to 1.00, indicating convergence, and ESS values were >1,000, indicating many uncorrelated draws were obtained (Muth et al. 2018).

**Fig. 3.**
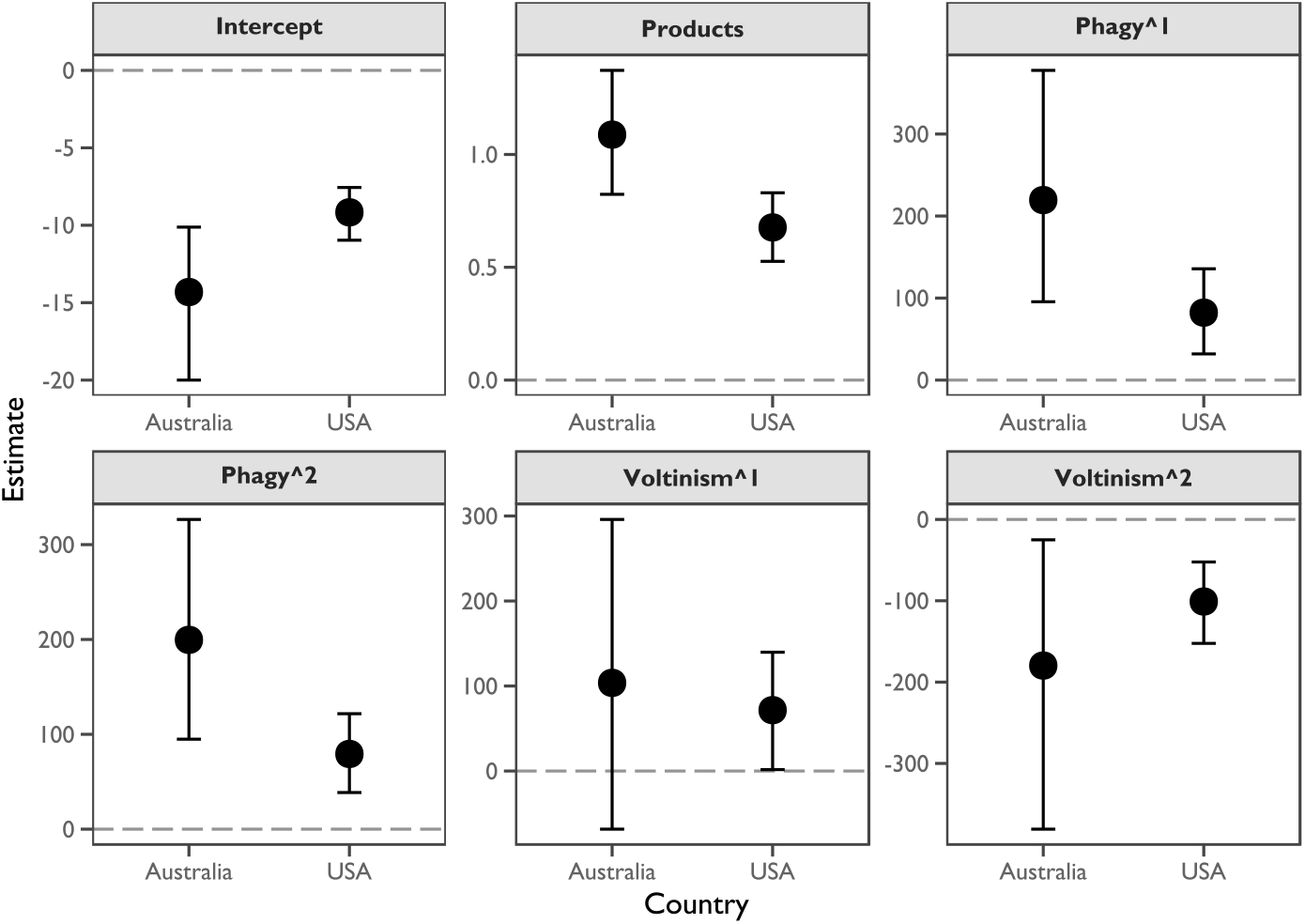
Intercept and partial regression coefficient estimates in models of resistance. Each panel represents a modelled term, with the *x*-axis representing the country, and the *y*-axis representing the coefficient estimate, with Bayesian 95% credible intervals. The dashed line demarcates an estimate of zero.

Our ‘full’ model including pesticide product registrations and ecological traits were substantially more likely (log-Bayes factor *≫* 1.1) to explain resistance status of a species to different chemical MoAs relative to ‘null’ model excluding these factors. For the USA species set, the ‘full’ model had an estimated log-Bayes factors of 88.24 and a Bayesian *R*^2^ of 0.33. For the Australian species, the ‘full’ model had an estimated log-Bayes factors of 83.04 and a Bayesian *R*^2^ of 0.45. Additionally, the ‘products’ model had a log-Bayes factor of 45.85 for the USA species set, with a Bayesian *R*^2^ of 0.31. For the Australian species set, the ‘products’ model had a log-Bayes factor of 29.85 and a Bayesian *R*^2^ of 0.34. Therefore, models fitting only pesticide products as a fixed effect were substantially more likely to explain resistance status of a species to different chemical MoAs relative to the ‘null’ model.

### Model generality

*K*-fold cross-validation was used to test the predictive power of the ‘full’ and ‘products’ models using 100 training-testing species set partitions. In each partition, the training species were used to construct the model, predictions were made on the testing species, and an AUC score was calculated. Predictive power was similar between models and countries (Fig. 4). For the USA species set, the mean AUC scores were 0.81 (±0.004 SEM) for the ‘full’ model and 0.80 (±0.003 SEM) for the ‘products’ model. For the Australian species set, the mean AUC scores were 0.82 (±0.007 SEM) for the ‘full’ model and 0.79 (±0.006 SEM) for the ‘products’ model. AUC scores were obtained using only the estimated fixed effect coefficients and not the estimated random intercepts for taxonomy or chemical MoA group. Hence, these *K*-fold cross-validation analyses show that resistance can be predicted well without controlling for ecology, taxonomy, and chemical MoA group, because species with resistance to a chemical MoA group tend to have greater product registrations than those susceptible to that chemical MoA group.

**Fig. 4.**
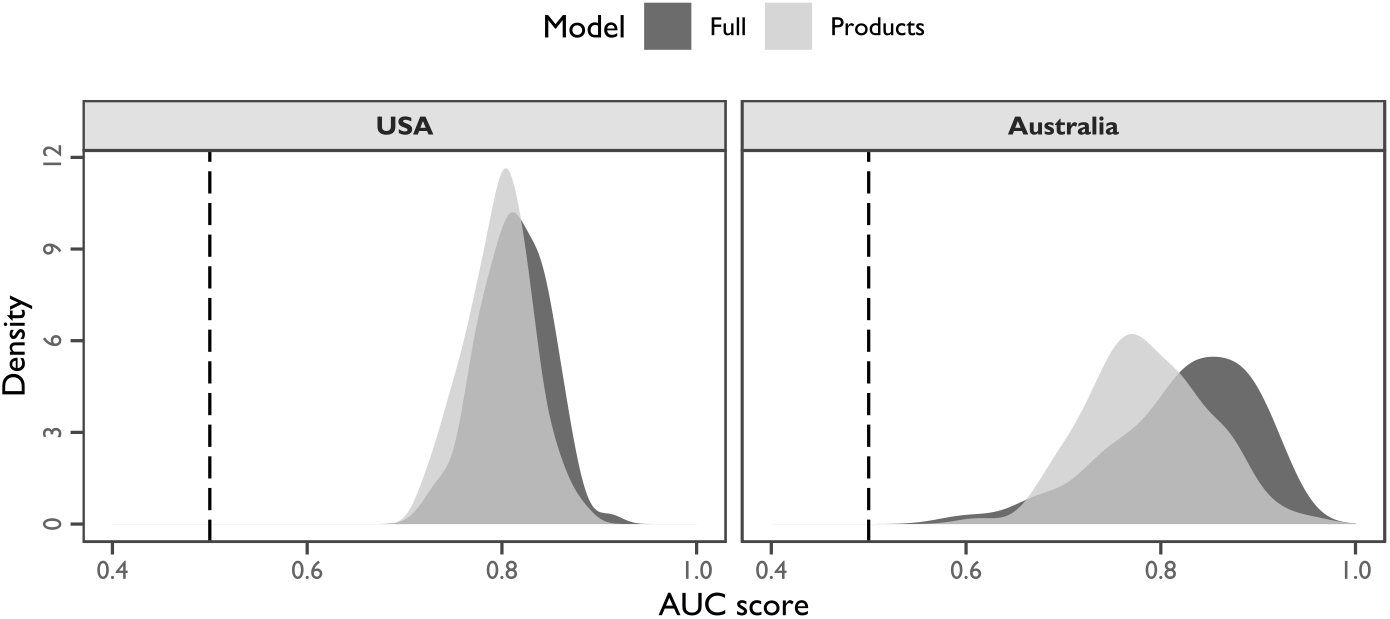
Comparisons of predictive power between models using AUC scores with 100 training–testing partitions. Curves depict the kernel density estimate, with area under the curve (AUC) equal to 1.00. Dark grey curves depict the distribution of AUC scores obtained from the ‘full’ model and light grey curves depict scores obtained from the ‘products’ model. The dashed lines depict an AUC score of 0.50, that is, an equal probability that the model assigns a higher score to resistant or susceptible species–chemical MoA observations. AUC scores approaching 1.00 indicate an increasing probability that resistance species–chemical MoA observations will be assigned higher scores than susceptible observations. Panels contain results from the USA and the Australian species sets separately.

## DISCUSSION

Identifying key predictors of pesticide resistance and pinpointing high-risk species increases capacity for making proactive, data-driven risk management decisions. Here, we extend our prior work by showing that pesticide products are not only positively associated with pesticide usage (Maino et al. 2023), but they are also positively associated with resistance status in arthropod pests. Although prior works have devised different frameworks for estimating resistance risk (Rotteveel et al. 1997; Moss et al. 2019) or modelling predictors of resistance (Rosenheim et al. 1996; Grimmer et al. 2015; Brevik et al. 2018; Hardy et al. 2018; Crossley et al. 2020), our study is the first to derive a covariate of resistance risk that is related to selection pressure, to the best of our knowledge. Our models of resistance risk predict the probability that a species is resistant to a chemical MoA (see also Rosenheim et al. 1996), but other studies have modelled the number of resistances in pests (Hardy et al. 2018), or the time for resistance to evolve (Grimmer et al. 2015; Brevik et al. 2018; Crossley et al. 2020). A unique aspect of our study is that we replicated our model of resistance risk in arthropod pests from two countries, the USA and Australia. This successful geographic replication allows us to place greater confidence in the generality and robustness of our conclusions. We believe pesticide product registrations hold great promise for predictive risk modelling and proactive resistance management, not only in arthropod pests, but in weeds and fungi as well, quantitatively linking species and potential selection pressures imposed by different chemical MoAs.

We hypothesize that the positive association between pesticide product registrations and resistance status arises from the accumulation of processes that generally increase the selection pressure on pest populations. Note that we do not imply to have estimated selection coefficients themselves, but instead have defined a covariate for modelling purposes that correlates with selection pressure. The risk of pesticide resistance in any pest is expected to be relative to the magnitude and frequency of selection (Georghiou and Taylor 1977; Evans et al. 2016; Hicks et al. 2018; Hawkins et al. 2019). Additionally, for a chemical MoA to be extensively used, it must be accessible, affordable, and in demand (Sparks and Nauen 2015; Umina et al. 2019). Chemical databases may therefore capture trends in both the availability and application intensity of chemicals in the field, which translate into greater selection pressures in the field, and therefore risk of resistance evolution.

We see the application of pesticide product registrations to proactive management as a two-step procedure. First, as we have done here, a model fitting resistance status of species to chemical MoA groups needs to be constructed. Second, after fitting this model across species and chemical MoAs, species with high predicted risk scores for a chemical MoA but with no documented resistances can be identified as candidates for further inquiry. We illustrate such predicted risks for the top ten resistant pests (those with the most resistances to different chemical MoAs) in the USA (Fig. 5a) and Australian (Fig. 5b) species sets. We highlight the three species–chemical MoA combinations with highest predicted risks from an Australian context:

1. ***Tetranychus urticae*, the two-spotted mite, to avermectins (MoA group 6, ŷ = 0.69)**. *Tetranychus urticae* has resistance to 12 different chemical MoAs in Australia, and there are only six chemical MoAs for which *T. urticae* is susceptible. Within Australia, avermectins are widely used to control *T. urticae* in grain, vegetable, cotton, and fruit crops (Australian Pesticides and Veterinary Medicines Authority 2021; Dodds and Fearnley 2021; Grundy et al. 2021). Avermectins are the third most registered chemical MoA for *T. urticae* (following organophosphates and pyrethroids), so resistance to this chemical MoA would constitute a non-trivial loss of chemical control options. Field resistance of *T. urticae* to avermectins has already been reported in other parts of the world, involving metabolic mechanisms (Stumpf and Nauen 2002), and reported laboratory-evolved resistance has been underpinned by target-site mutations (Kwon et al. 2010; Dermauw et al. 2012). Our results indicate Australian *T. urticae* populations have a moderate risk of evolving resistance to avermectins, highlighting the value of screening field populations to assess the status of local resistance to this chemical group.
2. ***Epiphyas postvittana*, light brown apple moth, to pyrethroids (MoA group 3A, ŷ = 0.47)**. Reported resistances in this pest are to older chemical groups (*i.e*., carbamates, organophosphates and organochlorines) from the 1960s through to the 1980s (Kerr 1964; Mota-Sanchez and Wise 2021). There are many chemical options for *E. postvittana* control, but pyrethroids are the third most registered chemical MoA for *E. postvittana* in Australia (following organophosphates and carbamates) and are used for control of this pest in fruit, vegetable, and ornamental crops (Australian Pesticides and Veterinary Medicines Authority 2021). The predicted risk of pyrethroid resistance evolution is moderate, although judicious chemical rotations and use of non-chemical management options available for *E. postvittana* control (Suckling and Brockerhoff 2010) could help mitigate this risk.
3. ***Helicoverpa armigera*, corn earworm, to amitraz (MoA group 19, ŷ = 0.46)**. Amitraz is primarily used in the Australian cotton industry to manage *H. armigera*, with up to four applications often recommended per growing season (Grundy et al. 2021). Amitraz is the only chemical MoA group currently registered against *H. armigera* for which no current resistance data is available in Australia (Grundy et al. 2021), although it was presumably still susceptible in the early 2000s (Gunning 2002). *Helicoverpa armigera* is already resistant to eight chemical MoA groups in Australia, many of which are widely distributed throughout the country (Thia et al. 2021). Our results suggest a moderate risk of amitraz resistance, even though no reports of resistance to this chemical group have been reported for *H. armigera* globally (Mota-Sanchez and Wise 2021). Although it is unclear how this predicted risk may translate into actual risk in the field, it would be useful to collect baseline data for amitraz susceptibility in Australia.

**Fig. 5.**
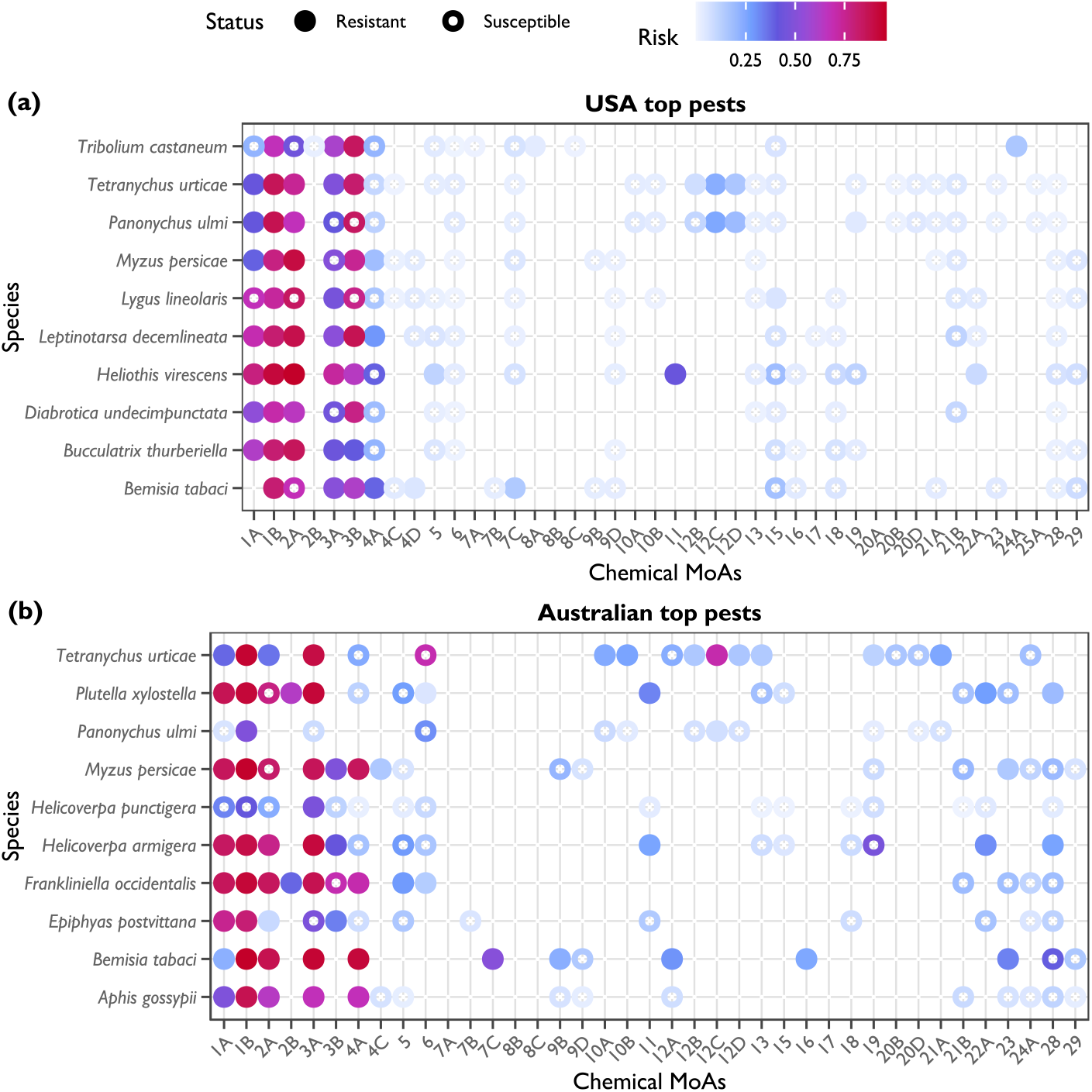
Predicted risks for the ten most resistant agricultural pest species in the USA and Australia. Closed circles represent species–MoA combinations that exhibit resistance, whereas open circles represent species–MoA combinations that are susceptible. Circles are coloured by the magnitude of predicted risk, with low risk in pale blue (0.00) to high risk in red (1.00). Only risks for chemical MoAs registered for each pest species are shown. (a) The USA top ten pest species with most cases of resistance (field evolved resistances reported in the APRD). (b) The Australian top ten pest species with most cases of resistance (field evolved resistances reported in the APRD and literature search).

Consistency in the effect of pesticide product registrations in both the USA and Australian models of resistance status is a noteworthy outcome of our study. Such a finding mightbe partially due to shared characteristics of the cropping industries and agro-ecosystems of these countries. For example, the taxonomic composition of agricultural arthropod pests was similar between countries; the orders Lepidoptera and Hemiptera had the most registered pest species, followed by Coleoptera and mites from the Trombifidormes (Fig. 1). However, at the species level, both countries have different pest communities. In both the USA and Australia, cotton and grains are the highest contributing cropping commodities to total earnings from agricultural production. In the USA, corn, soybeans, wheat, and cotton make up 15%, 11%, 3% and 2% of the total earnings (respectively), and in Australia, wheat, barley, canola, and cotton make up 13%, 5%, 4%, 3%

(respectively) (Howden and Zammit 2019). Whilst the effect size of pesticide product registrations on resistance status was similar between the USA and Australia, application of this predictor into other geographic regions will require construction of region-specific datasets and models of resistance status. We recognize that the use of pesticide product registrations as a predictor of resistance status will be greatest in geographic regions where positive correlations to field usage exist. In developing and lower-income countries, pesticide usage can be less intensely regulated (Ecobichon 2001; Schreinemachers and Tipraqsa 2012), which may decouple the association between pesticide usage and product registrations. Replicating the use of pesticide product registrations in agro-ecosystems of different countries represents an important direction for future work, as does the extension of our framework to other agricultural pests, such as weeds and fungi, by compiling similar datasets of specific resistances and product registrations to herbicides and fungicides.

Whilst pesticide usage is necessary to impose selection, the ecology of pests play an important role in the rate at which resistance evolves. Our work demonstrates resistance evolution in Australian agricultural pests exhibits similar curvilinear relationships with phagy and voltinism as that observed in the USA (Rosenheim and Tabashnik 1991; Rosenheim et al. 1996; Hardy et al. 2018; Crossley et al. 2020). There was a striking consistency in both the direction and magnitude of effect in partial regression coefficients between the pest species sets from the USA and Australia (Fig 3). The positive second order effect of phagy suggests that pest species with an intermediate host plant breadth have the lowest risk of resistance, and that narrow and wide host plant breadth is associated with greatest resistance risk. For more specialized pests with narrow host plant breadth, a greater proportion of the population is constrained to treated crops and exposed to selection (Georghiou and Taylor 1986). For more generalist pests with wide host plant breadth, a greater arsenal of biochemical machinery used to metabolize plant phytochemicals are available to detoxify pesticides (Dermauw et al. 2018; Rane et al. 2019). The negative second order effect of voltinism suggests that pest species with an intermediate number of generations per year have the greatest risk of resistance, and that few and many generations per year is associated with a lower risk of resistance. The curvilinear relationship between voltinism and resistance risk has a less clear interpretation. In some cases, voltinism has been described as a nuisance parameter (Crossley et al. 2020). However, the number of generations per year may modulate the effect of other operational, ecological, and population genetic factors on resistance evolution (Georghiou and Taylor 1977; Rosenheim and Tabashnik 1990). Regardless of the underlying mechanism, our analyses show that phagy and voltinism may be truly general predictors of resistance evolution risk in agro-ecosystems.

The predictive framework developed here is not without limitations, including: (i) legacy effects captured in pesticide product registration and resistance status datasets; (ii) ephemeral resistances; (iii) lag between the evolution of resistance and the accumulation of pesticide product registrations; (iv) methodological effects of assigning pesticide product registrations and model fitting; and (v) the coarse scale of our predictions.

i. Legacy effects are associated with analysis of temporally accumulated data. For pesticide registration data, we summed the cumulative number of products across years, and for resistance status, we scored pests as resistant to a chemical MoA if there was at least one record, irrespective of when it was reported. Some MoA groups are no longer used in many countries due to their negative health and ecotoxicology effects, such as organochlorines and DDTs, or because resistance is so commonplace they are no longer efficacious, such as pyrethroids for *Frankliniella occidentalis* (western flower thrips) control in Australia (Herron and Gullick 2001).
ii. Ephemeral, transient resistances also contribute toward legacy effects. Some species periodically exhibited high levels of resistance, presumably in response to intense short-term selection pressure, which later disappears. In Australia, two such examples include *Therioaphis trifolii* (spotted alfalfa aphid) resistance to carbamates and organophosphates (Holtkamp et al. 1992) and *Helicoverpa punctigera* (native budworm) resistance to pyrethroids (Gunning et al. 1997).
iii. Lag effects may occur if pests evolve resistance to a chemical MoA before pesticide product registrations accumulate to a notably “at risk” level. We attempted to account for this by fitting random intercepts for taxonomy and chemical MoA. But we also found that models including pesticide product registrations alone (no ecological predictors) could predict resistance status (Fig. 4). Hence, despite being an approximate measure of pesticide usage, we believe that pesticide product registrations sufficiently covary with resistance status to be a useful predictor of resistance risk in agricultural pests that are currently susceptible.
iv. Assigning pesticide product registrations required decisions on how to assign taxonomically ambiguous pest names and chemical mixtures. In both the USA and Australian chemical databases, many pesticide products were registered against pest common names that could not be assigned to the species level (genus or higher). To overcome this issue, we assigned pesticide product registrations to all species nested within the relevant taxonomic rank (up to the family level), making all species within a taxonomic rank more similar. In our analysis, chemical mixtures contributed counts to each unique chemical MoA present in a pesticide product. This effectively assumes that chemical mixtures apply selection for all constituent chemical MoAs and that their effect is equivalent to pesticide products comprising a single chemical MoA. Treating pesticide product registrations in this way was necessary to automate data curation, exploit as much information as possible from chemical databases, and avoid fitting complicated models; however it does not take into account that some active ingredients in a mixture may be ineffective against certain pests. These methodological effects are expected to add noise to predictions and reduce power. For example, a predicted risk of neonicotinoid resistance (MoA group 4A) for *H. armigera* (albeit a low risk) is unusual given neonicotinoids have low physiological activity against many Lepidopterans relative to other pests (reviewed in Elbert et al. 1998). This observation is being driven by registrations of mixtures containing neonicotinoids with other active ingredients that target *H. armigera*, such as avermectins (MoA group 6) and diamides (MoA group 28).
v. Spatially and temporally patchy resistances are somewhat misaligned with our models, which have coarse spatio-temporal resolution and consider all observations as contemporaneous. Whilst the risk predictions produced by our models are useful for identifying candidate species for proactive management at a region-wide level, they lack the fine-scale resolution necessary to guide action at local levels. Although localized models can be successfully developed, they are almost always constrained to single species (Feng et al. 2010; Ives et al. 2017; Maino et al. 2018; Ma et al. 2021). Research and data management infrastructures that capture fine-scale chemical usages patterns and spatio-temporal variability in pesticide tolerance phenotypes across many pest species are needed (Thia et al. 2021). Such data is necessary to generate flexible multi-species predictive models at scales relevant for localized management.

## CONCLUSIONS

Developing analytical methods that delimit ‘high-risk’ versus ‘low-risk’ species can guide future resistance monitoring efforts and facilitate decisions on resource allocation. We detail the novel use of pesticide product registrations as a predictor of resistance in agricultural arthropod pests from the USA and Australia. Predicted risks of resistance can help pinpoint when the intensity and duration of chemical selection in the field may warrant attention for specific pest and chemical MoA group combinations. However, we note that it will be important to interpret these risk predictions in the context of pest biology, the broader agro-ecosystem, and expert opinion when making management decisions. Nonetheless, our work should aid the development of proactive pest management programs in agro-ecosystems.

## ACKNOWLEDGEMENTS

We thank David Mota-Sanchez and John Wise for their permission to use the Arthropod Pesticide Resistance Database records in our study. We acknowledge Kaya Moore, Candida Wong, Courtney Brown and Safieh Soleimannejad for helping curate taxonomic information and ecological trait data. Computational resources were provided by The University of Melbourne’s Research Computing Services and the Petascale Campus Initiative. We thank two anonymous reviewers for their constructive comments on this manuscript.

## DECLARATIONS

### Ethical Approval

Not applicable

### Consent for publication

Not applicable.

### Authors’ contributions

PU and AH conceived the project and secured funding. JT and JM designed the study and compiled the pesticide registration data. AK led the compilation of trait data. JT led the statistical analysis and drafted the original manuscript. All authors contributed revisions and approved the manuscript.

### Availability of data and materials

All data and scripts required to reproduce the analyses presented in this manuscript have been uploaded to Dryad (Thia 2022), DOI: 10.5061/dryad.7pvmcvdwc.

## LISTS OF KEYWORDS FOR DATABASE QUERIES AND LITERATURE SEARCHES

### List S1. Key words used to filter the EPA database of chemical registration in the USA

- BAIT APPLICATION
- BARK TREATMENT
- DIP TREATMENT
- DORMANT APPLICATION
- FOLIAR TREATMENT
- FORAGE
- GRAIN
- GREENHOUSE
- NURSERY
- PASTURE
- POSTHARVEST
- ROOT
- SEED
- SEEDLING
- SEED CROP
- SEED TREATMENT
- SOIL FUMIGATION
- SOIL TREATMENT
- WATER TREATMENT

### List S2. Key words used to search for phagy values in Google Scholar

- food plant
- host
- hosts
- host plant
- host plants
- phagy

### List S3. Key words used to search for voltinism values in Google Scholar

- biology
- ecology
- generation per year
- generations per year
- life cycle
- life history
- multivoltine
- phenology
- reproductive rate
- univoltine
- voltinism

## TABLES AND FIGURES

**Table S1.**
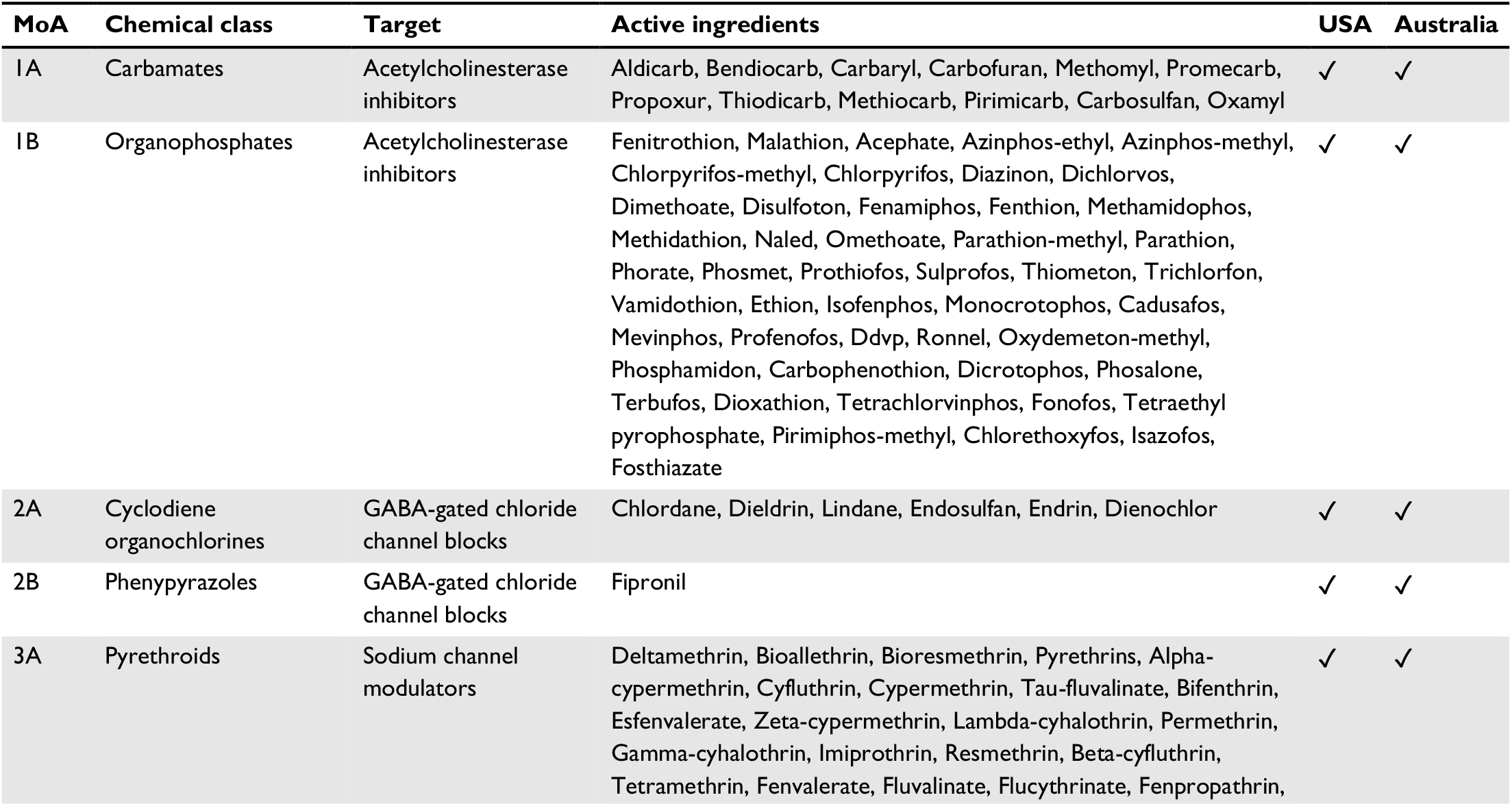
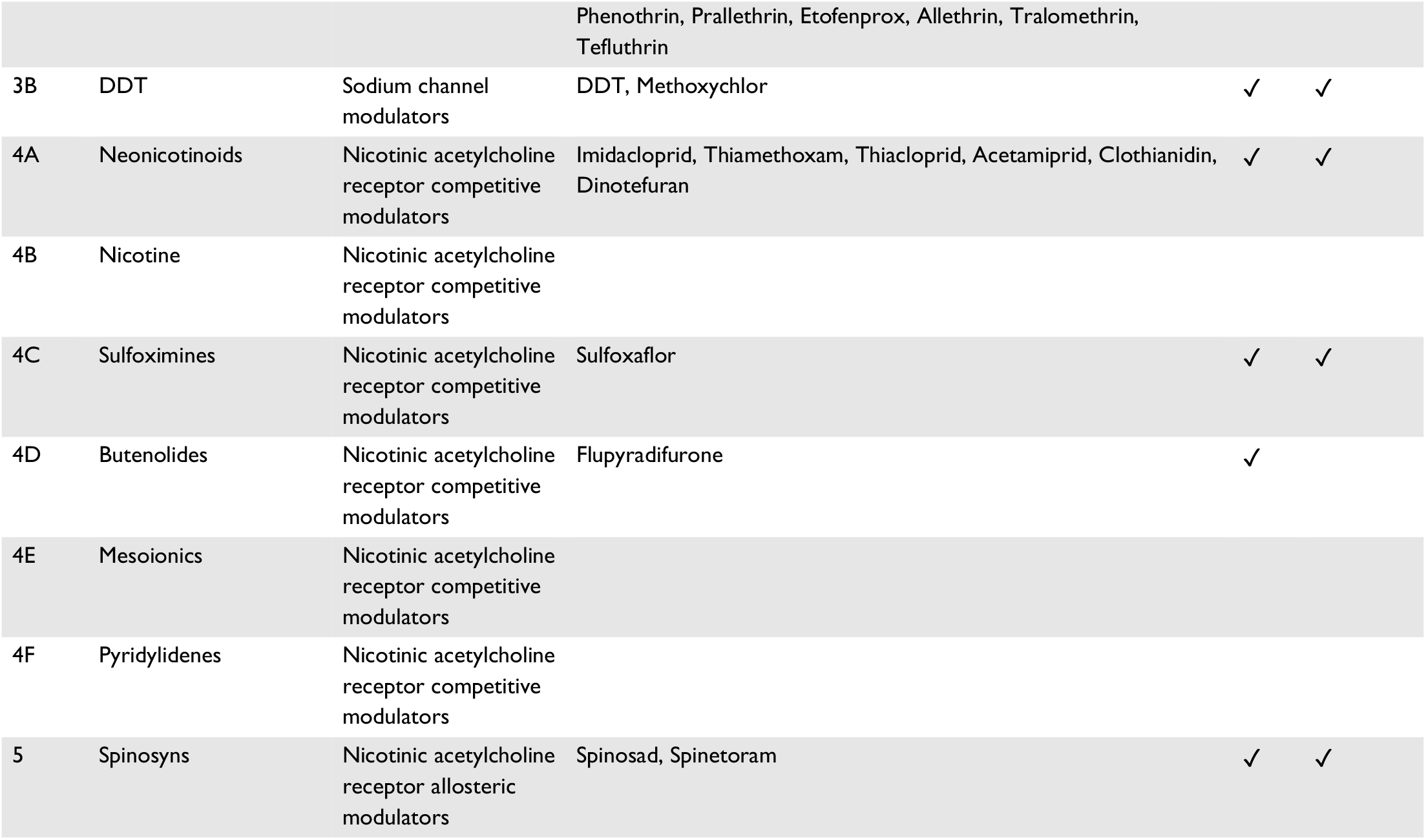
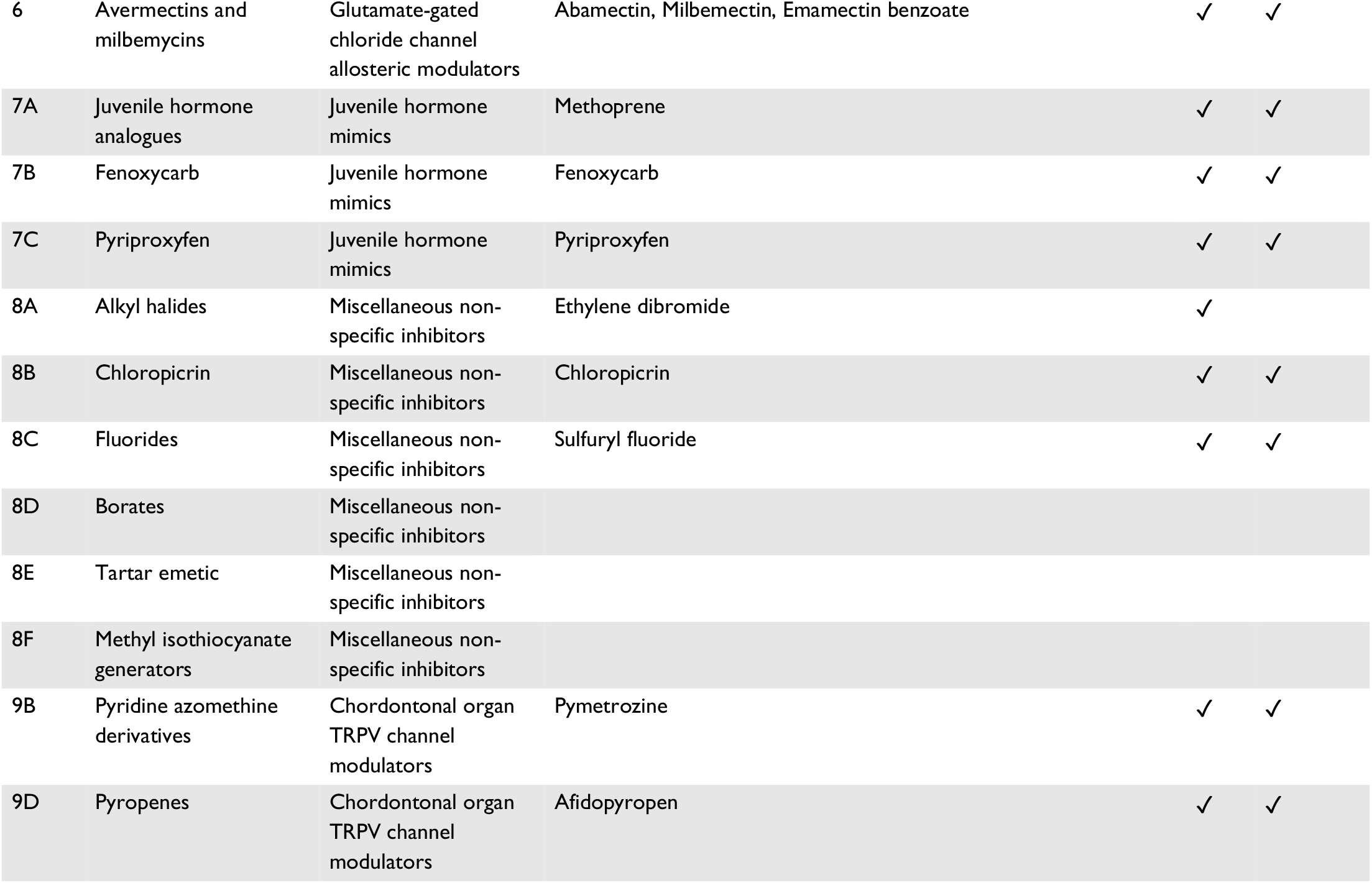
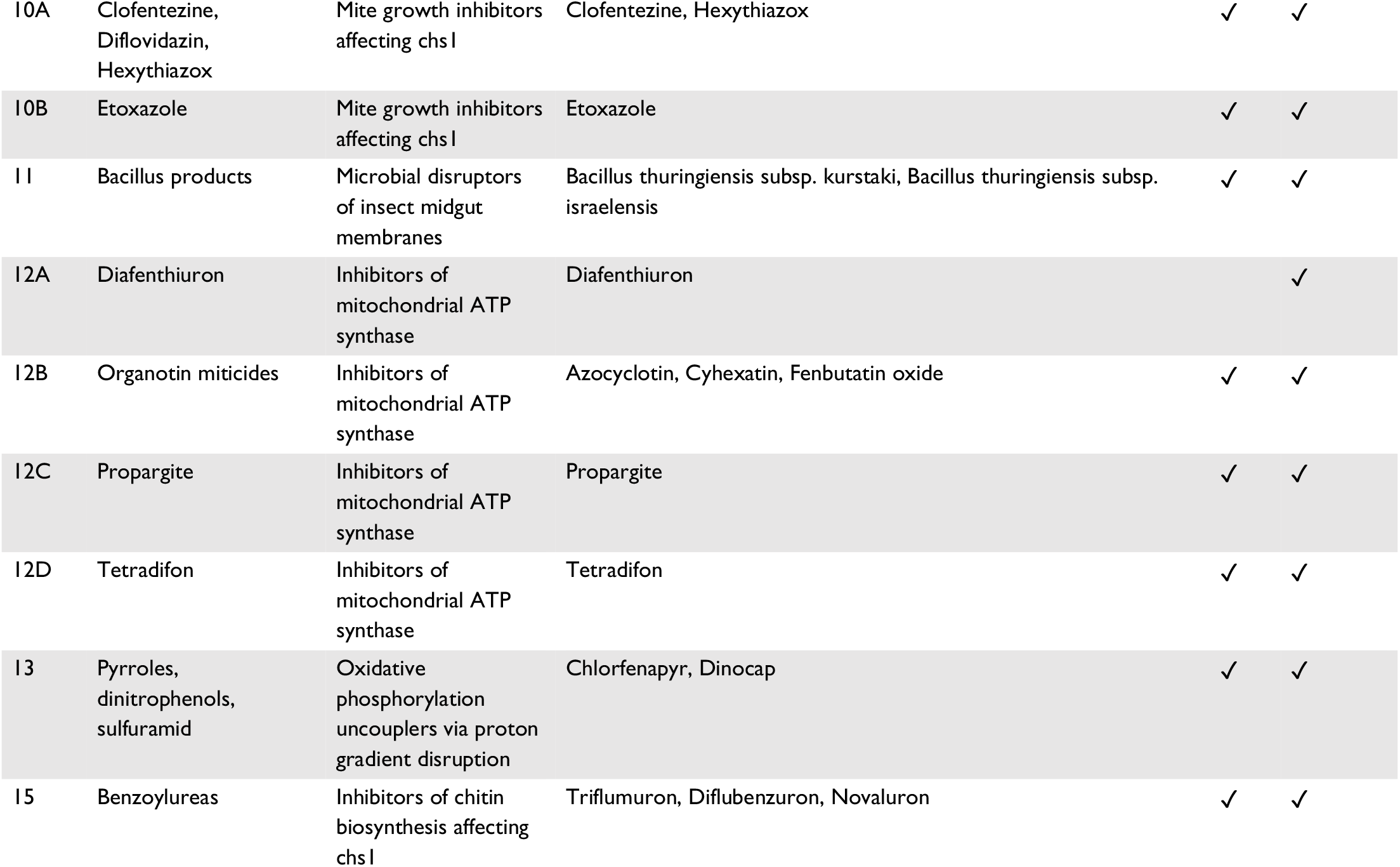
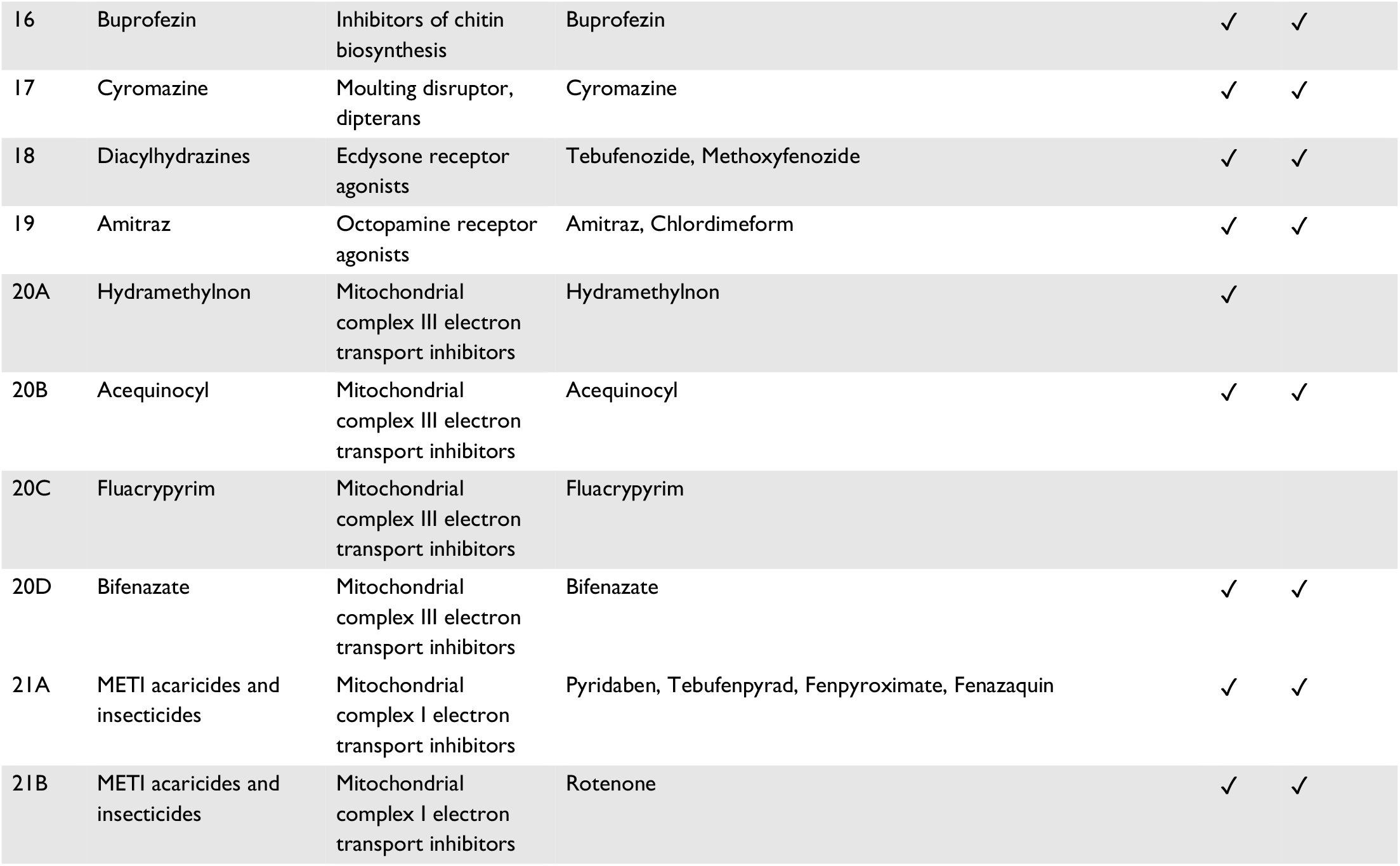
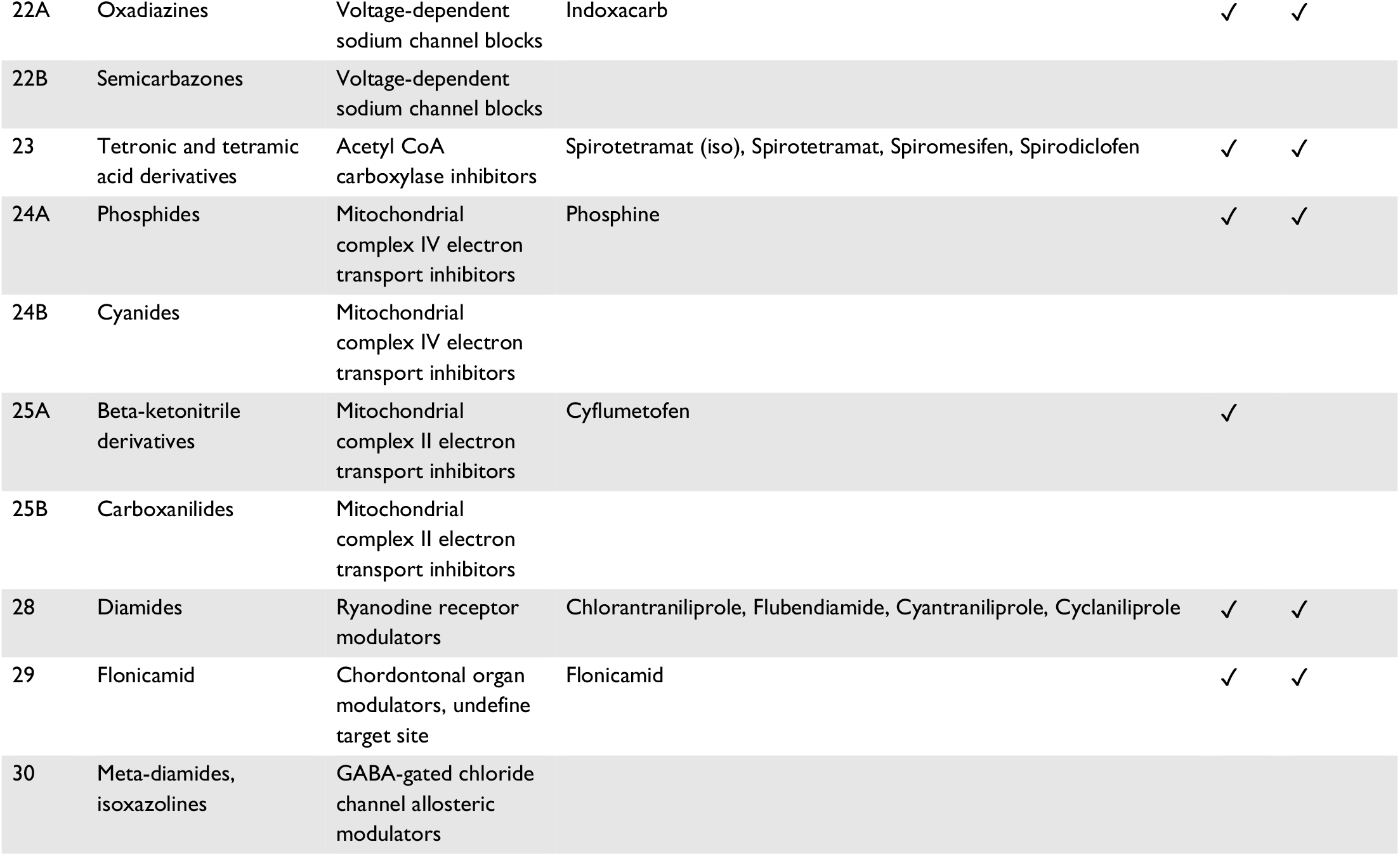
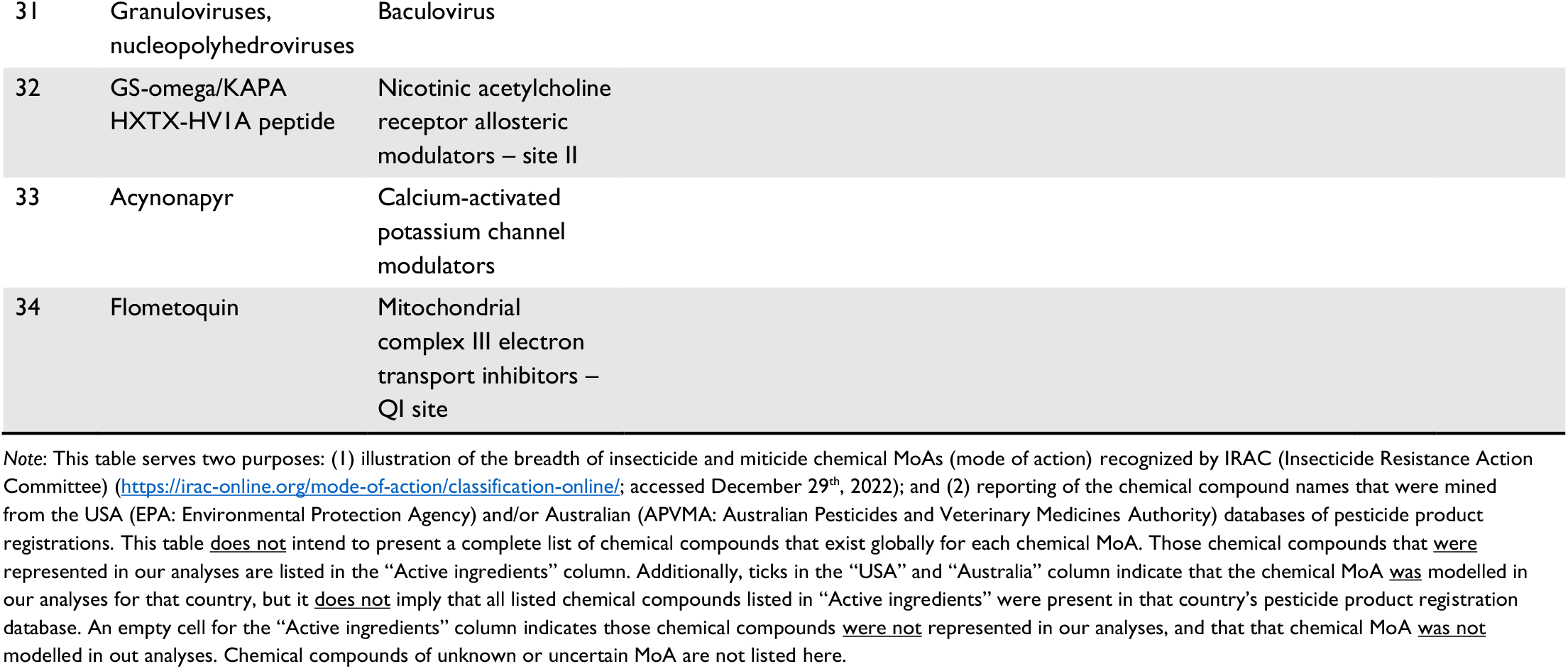
Chemical MoA groups and active ingredients included in this study for the USA and Australia.

**Fig. S1.**
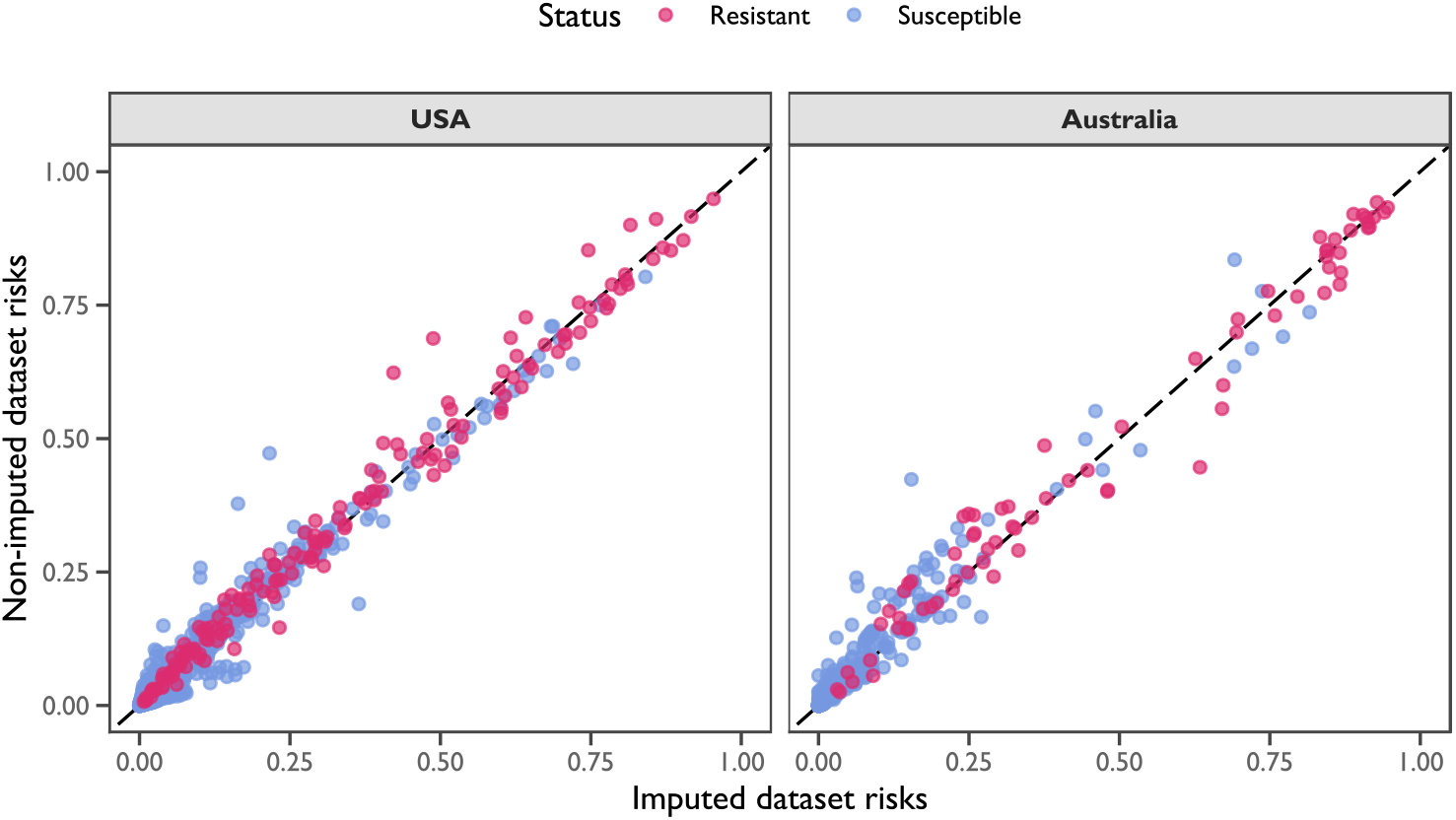
Comparisons of predicted resistance risks from the ‘full’ model using datasets with imputed and non-imputed ecological traits. Points represent predicted risks for species–chemical MoA combinations, coloured by resistance status: pink for resistant, and blue for susceptible. The dashed line demarcates the 1:1 association. Panels contain predictions for the USA (left) and Australian (right) species sets.

